# The influence of talus size and shape on *in vivo* talocrural hopping kinematics

**DOI:** 10.1101/2024.02.15.580586

**Authors:** Anja-Verena Behling, Luke Kelly, Lauren Welte, Michael J Rainbow

## Abstract

Talus implants often come in standard sizes and shapes; however, humans vary in their bone size and shape. Consequently, patient-specific implants are becoming available. Understanding how shape changes alter function in a healthy cohort may help designers determine how much specificity is required in talocrural implants.

Nine participants (5 females) hopped on one leg while biplanar video radiography and force plate data were collected. 3D bone models were created from computed tomography scans. Helical axes of motion were calculated for the talus relative to the tibia (rotation axes) and a cylinder was fit through the talar dome (morphological axis). Bland-Altman plots and spatial angles tested whether the rotation and morphological axes agree. A shape model of 36 (15 females) participants was established and a cylinder fit was morphed through the range of ±3 standard deviations.

The rotation and morphological axes largely agree regarding their orientation and location during hopping. The morphological axis consistently overestimates the orientation-component in anterior-posterior direction. Some shape components affect talar dome orientation and curvature independent of size. This suggests that besides size, the shape of the talar dome might affect the movement pattern during locomotion. Our findings are important to inform talocrural joint arthroplasty design.

## 1. Introduction

Ankle arthroplasties aim to mimic talus anatomy and function; however, guidance on their design is limited because there is minimal information linking *in vivo* talocrural joint function to talus morphology. Shape studies have shown that talus bone size and shape alter talar dome curvature and orientation [1–3] and kinematic studies have shown that the talocrural joint displays variable out-of-sagittal-plane kinematics, violating the common assumption [4,5] that the talocrural joint behaves like a hinge [6–8]. However, the complex changes in the size and shape of the talus have not been directly linked to changes in talocrural joint function. Fortunately, recent advancements in biplanar video radiography (BVR) can address this gap [9–11].

Size is the largest source of overall shape variation in the human talus [12–14]. In other joint complexes, such as the wrist, size explains the distal and proximal location of the axis of rotation [15,16]. Based on observations at the wrist, we might expect to observe that smaller tali would have a more superiorly located rotation axis than larger tali (e.g., closer to the tibia). Understanding the curvature of the talar dome is important because it might affect moment arms, ligament and tendon resting lengths, and articular contact stress [17,18]. However, the curvature of the talar dome may also vary across individuals independent of size. In other words, the shape of the talus may also influence the location of the axis. Since a cylinder fit to the talar dome approximates its curvature [5,19], a morphology-based cylinder axis may serve as a reasonable proxy to describe changes associated with variations in size and shape due to changes in talar dome curvature.

While the talar dome’s curvature due to size or shape variation may explain the superior/inferior location of the rotation axis and, talocrural motion may be more difficult to capture by a single morphologically based cylinder axis. If the talocrural joint behaves as a hinge (not necessarily only positioned in the sagittal plane) and the mediolaterally symmetric changes in sagittal curvature are the primary drivers of talocrural motion, we would expect the orientation of the cylinder axis and rotation axis to be in close agreement. It is unclear if the cylinder axis captures mediolateral asymmetric variation and curvature changes about the transverse and coronal planes. Understanding how well the orientation of the rotation axis and cylinder axis agree may provide insight into the aspects of talar dome morphology that are important to talocrural function.

This study aims to quantify whether the morphological axis based on a cylinder fit captures the talus’s rotation axis and to determine the morphological determinants (size and shape) of these axes. Here, we use a novel approach that links *in vivo* kinematics from BVR to geometric morphometrics. We chose hopping as our kinematic task because it maximizes the range of motion at the talocrural joint. Based on the literature, we expect that there will be good agreement across participants in the morphological and rotation axes. We also predict that the axes’ superior/inferior location will vary as a function of talus size, where a larger talus corresponds to a further inferiorly positioned axis location relative to the tibia. Finally, we determine which shape features, independent of size, affect the axes.

## 2. Methodology

### 2.1. Overview and Participants

After institutional review board approval from Queen’s University and written informed consent was obtained (MECH-061-17), nine healthy participants (5 females; mean ± std, mass 71.3 ± 14.5 kg; height 171.4 ± 10.7 cm) with no history of lower limb injury hopped barefoot on their right leg while matching their hopping frequency to a metronome at 156 bpm. BVR (125 Hz, range of 70–80 kV, 100–125 mA) and synchronized ground reaction forces (1125 Hz, AMTI Optima, AMIT, Watertown, MA) were collected for three hops during each sustained hopping trial.

### 2.2 Data Processing: Kinematics Derived from BVR

A computed tomography scan (120 kV, 60mA, model: Lightspeed 16, n = 9, Revolution H.D., General Electric Medical Systems, USA) was obtained from the right foot while participants lay in a prone position with their feet plantar flexed (average resolution: 0.356 × 0.356 × 0.625 mm). Tibia and talus bones were segmented (Mimics 24.0, Materialise, Leuven, Belgium), yielding bone surface meshes and partial volumes (an image volume of the isolated bone of interest) [20].

We used an established processing workflow to track footbones with BVR [21]. Briefly, the high-speed cameras were calibrated using a custom calibration cube, and the images were undistorted [22] using custom software (XMALab, Brown University, USA) [23]. The translation and orientation of the tibia and talus were measured by matching digitally reconstructed radiographs generated from the partial volumes to two calibrated radiographs for each frame of data. Tracking was done in Autoscoper (Brown University, USA) [20].

Gait events were defined using a 15N threshold in the vertical ground reaction force to determine touchdown and take-off. The transition point between the landing and push-off phase was determined at the active peak (maximal vertical ground reaction force after impact peak). Landing describes the phase from touchdown to the transition point and push-off from the transition point to take off.

The coordinate systems for the tibia and talus were shape-based, and their primary axes were determined by a cylinder fit through the distal tibia [18,19] and talar dome surfaces [26], respectively. For the tibia, the superiorly pointing axis is based on the cylinder orientation from a cylinder fit through the tibia shaft, and the anterior-posterior axis is obtained as the cross-product of the superior and primary axis. For the talus, A sphere is fitted through the talar head, another sphere through the talus/s calcaneal facet and the subtalar axis runs through the centroid of both spheres. The superior axis is determined by the cross-product of the primary axis and the subtalar axis. Lastly, the anterior-posterior axis is re-calculated via the cross-product of the superior and primary axis. The origins are in the centroid of each bone. The coordinate system axes were re-labeled such that the *x*-, *y*- and *z*-axes approximate dorsiflexion, inversion, and adduction, respectively, with reference to the right foot and right-hand rule [25,26].

We calculated finite helical axes for the talocrural joint resolved in the tibia coordinate system between two time points [27] (Supplement Figure S1). This results in two hopping phases: the landing phase (from touchdown to transition point) and the push-off phase (from the transition point to toe-off). Each hopping phase corresponds to one helical axis. A helical axis is defined by the rotation about and translation along an axis, termed the rotation axis [15,16]. The axis is also located by a point in space that is dictated by the radius of curvature about which the talus is rotating.

### 2.3. Statistical Shape Model of Talus

To quantify differences in talus shapes, we created a shape atlas for the talus bone. Data from multiple, institutional review board approved studies at Queen’s University were pooled together and written informed consent from all participants was granted (MECH-060-17, MECH-061-17 & MECH-063-18), resulting in 36 tali (mean ± std; mass 73.1 ± 13.7 kg; height 171.5 ± 8.2 cm; 15F). A reference mesh was randomly selected after visually inspecting all bones for what we qualitatively determined to be average features. All tali were aligned to their inertial axes and followed by an iterative closest point and rigid coherent point drift (CPD) algorithm to ensure optimal alignment to the reference bone. The first non-rigid CPD was performed to register the meshes to the reference bone by selecting corresponding points on the bone meshes relative to the reference mesh while conserving the individual bone shape. A generalized Procrustes approach was performed to scale and further align the meshes in Procrustes space [28].

Afterward, a new reference mesh was calculated as the average mesh across all Procrustes meshes (arithmetic mean). Subsequently, a second non-rigid CPD was performed to register all Procrustes meshes to the new reference bone. The corresponding meshes were used as input for principal component analysis (shape-PCA). An overview of the steps can be found in the supplementary material (Figure S2). To separate the effects of shape and size on rotation axis’s location and orientation, all further shape analyses (such as the cylinder fit) are carried out on the shape-PCA, where size has already been removed. For each principal component (PC) explaining the modes of variation within the PCA, the mean bone and bone deviations across ± three standard deviations (std) were reconstructed and visualized.

We calculated talus size using centroid size, which is the square root of the sum of the squared distances of all points of the bone mesh from their centroid, to understand the importance of bone size on rotation axes’ location and orientation.

### 2.4 Cylinder Fit Through Talar Dome

A talar dome template region was manually segmented for the new mean bone of the statistical shape model using Geomagic Wrap 2021.2.2. (3D Systems GmbH, Germany), which resulted in a 3D mesh of 180 points. Non-rigid CPD was used to determine the corresponding points on all participants’ talar domes based on the template region, as well as on the shape morphs. The corresponding talar dome points were then used as the input to fit a point cloud to a cylinder and further optimized using a least square cylinder fit method (Version R2021b, Mathworks, USA; Figure 1). The orientation and location of the cylinder (morphological axis) in the tibia coordinate system were assessed for each hopping participant (n = 9). To calculate the location of the morphological and rotation axis, the tibia coordinate system’s origin was moved to the tibial dome along the superior axis of the coordinate system. The reason for this was that the location of the original tibia coordinate system is influenced by how much tibia shaft was available in the CT scan and might therefore influence the outcome. The location of the morphological axis in superior-inferior and anterior-posterior directions was calculated relative to the origin of the re-adjusted tibia coordinate system origin. The cylinder’s curvature and therefore talar dome curvature is the inverse of its radius. Here, we report the radius because it allows us to make direct comparisons to the rotation axis.

**Figure 1:**
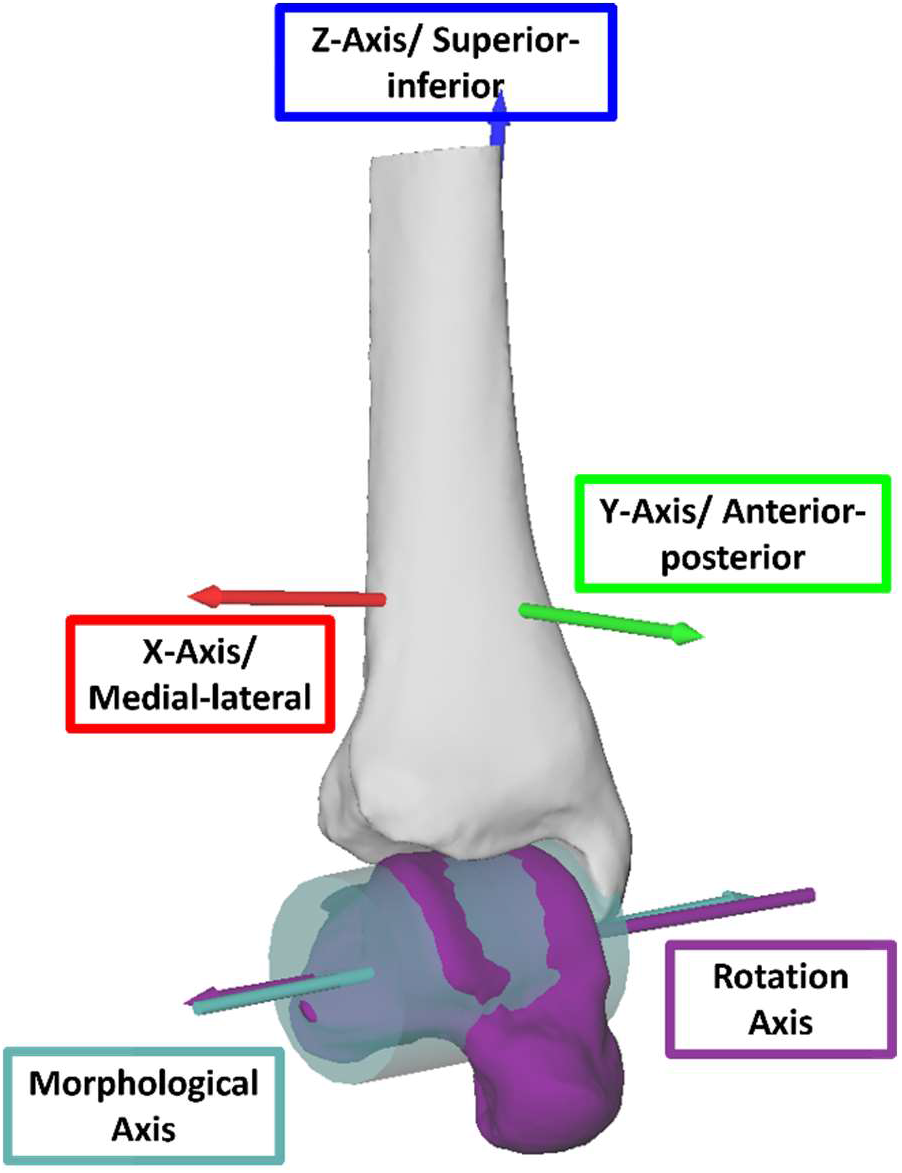
Cylinder fit to talar dome with the morphological axis (teal) and the mean average rotation axes for landing (purple) (n_Participant_ = 1; n_Trials_hopping_ = 3). Tibia coordinate system included (red, green, and blue arrow).

### 2.5 Analysis

Bland-Altman analysis determined the agreement between the morphological and rotation axes. We compared the orientation of the axes and report the spatial angles between the morphological and rotation axes. We also compared the location of the axes by projecting the morphological axis and rotation axis onto the sagittal plane of the tibia coordinate system.

To determine the effect of size on talar dome curvature, we performed a linear regression of the cylinder radius with talus size. To determine the effect of shape on talar dome curvature, we regressed the PC loadings of the shape-PCA on the cylinder radius. The level of significance was set to 0.05 unless indicated differently.

Initially, we applied a parallel analysis to determine the number of PCs significantly different from chance to determine the PCs of interest [29]. From the remaining PCs, each PC explaining more than 10% of the overall variance was visually inspected for changes in the shape of the talar dome, respectively.

## 3. Results

### 3.1. Agreement of Morphological & Rotation Axes

Agreement between the morphological and rotation axes was comparable for the landing and the push-off phases with average spatial angles of 9.6 ± 3.5° during landing and 9.0 ± 3.4° during push-off (Supplement Table S1). Hence, we only report the landing phase here (Figure 2); results for the push-off phase and Bland-Altman plots can be found in the Supplements (Supplement Figures S3-S6). The morphological and rotation axes agreed in the anterior-posterior and superior-inferior axis locations as there was no systematic bias, and the limits of agreement were within ± 2mm. The orientation of the morphological and rotation axes agreed best in the medial-lateral direction, as there was no systematic bias, and the limits of agreement were narrow (± 0.05). The morphological axis systematically overestimated the orientation in the anterior-posterior direction (bias = -0.1) relative to the rotation axes (schematic figure in Figure 2), and the limits of agreement were wider than in the medial-lateral direction (± 0.1). The orientation in the superior-inferior direction showed no systematic bias but reported the widest limits of agreement (± 0.2), indicating poorest agreement in this direction.

**Figure 2:**
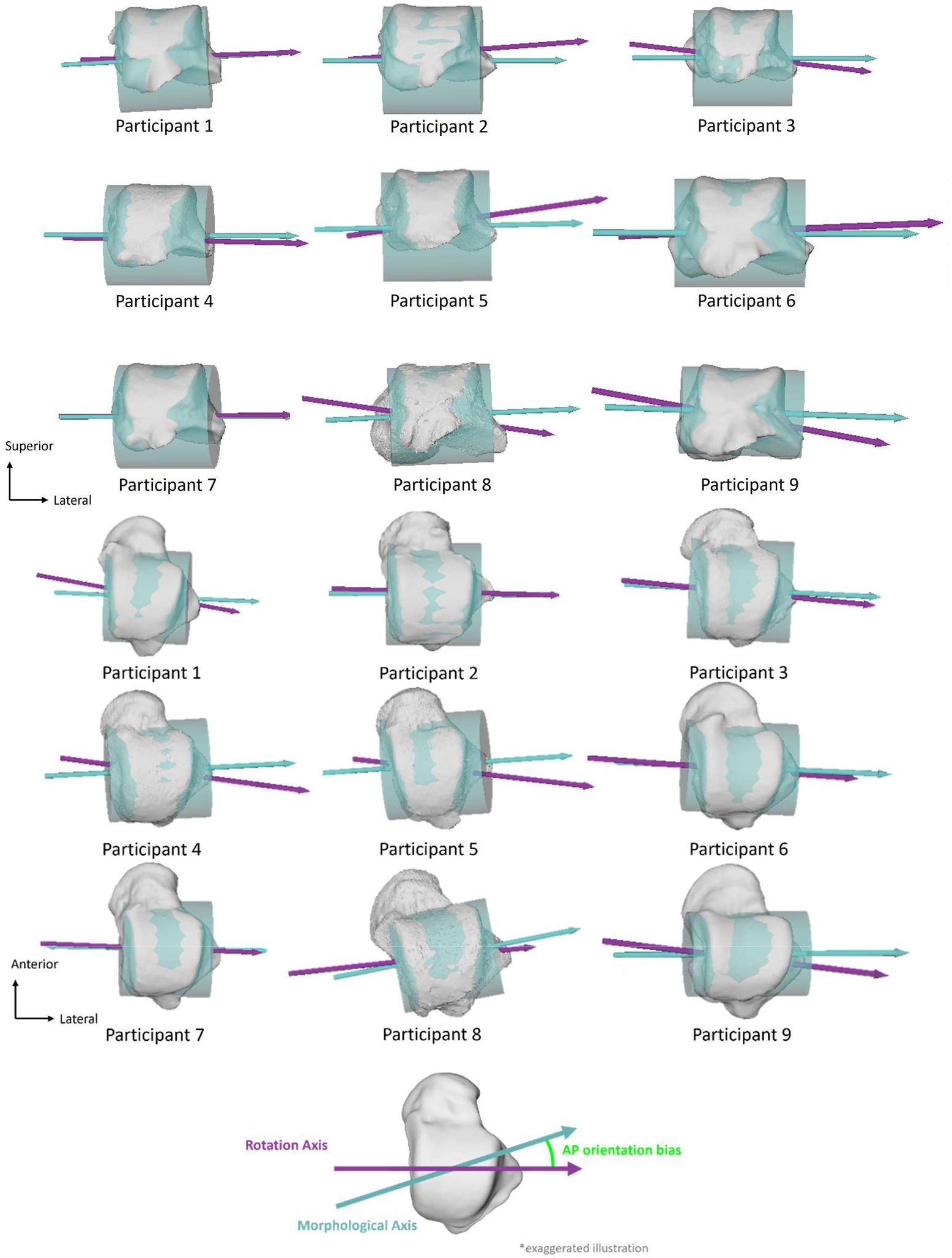
Orientation and location of rotation (average across all trials; purple color) and morphological (teal color) axes for each participant during landing. Schematic illustration of anterior-posterior bias of the morphological axis compared to rotation axis.

### 3.2. Effect of Size and Shape on Talus Dome Curvature

When looking at the influence of bone size on talar dome radius, larger tali were associated with larger cylinder radii (p = 0.002, r^2^ = 0.61, alpha = 0.05, Supplement Figure S3). The first three PCs of the shape-PCA explained 39% of the overall variance within the data set of scaled tali. Each PC explained more than 10% of the variance. None of the PCs were associated with centroid size (p ≥ 0.20; r^2^ ≤ 0.02; alpha = 0.05).

PC1 explains 16% of the overall variance in the data set and describes changes in talus posterior process prominence and talus head orientation and bulge. Within this PC across three standard deviations, alterations in talar dome metrics (cylinder radius and location) were within 1 mm (Figure 3). Changes in cylinder orientation and morphological axis orientation were below 0.01 in the medial-lateral and superior-inferior direction and small in the anterior-posterior direction (±0.1).

**Figure 3:**
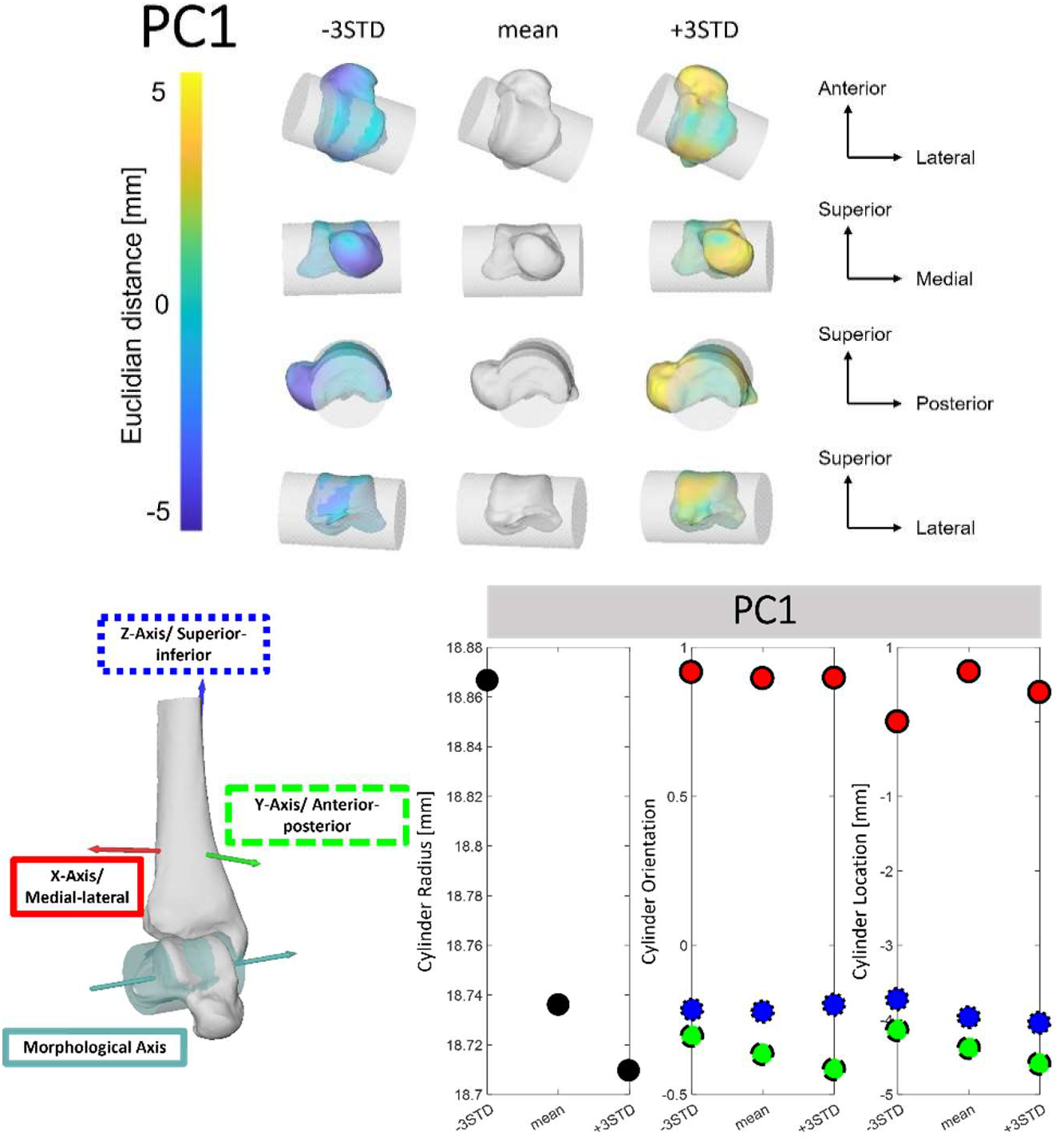
First PC of talus visualized with cylinder fit through talar dome ± 3 STD from the mean shape. Shape changes are color-coded according to the Euclidean distance in mm to the mean shape. Cylinder metrics for PC1 depicted across ± 3 STD for changes in cylinder radius (black), orientation and location with the three directions: medial-lateral (red and solid outline), anterior-posterior (green and dashed outline) and superior-inferior (blue and dotted outline). n_Participant_ = 36

PC2 explains 12% of the overall variance in talus shape. The main morphological variations can be seen in the posterior process prominence that “tucks under” the posterior talus region and lateral side of the talar head (shape changes indicated in dark blue and yellow across ±5mm, Figure 4). Consequently, the more curved the talar dome, the smaller the cylinder radius and the more superiorly and posteriorly located the morphological axis (associated here with +3STD changes). Changes in talar dome orientation occurred in the superior-inferior direction (internal-external rotation) but were small (±0.1) and even smaller in the medial-lateral direction.

**Figure 4:**
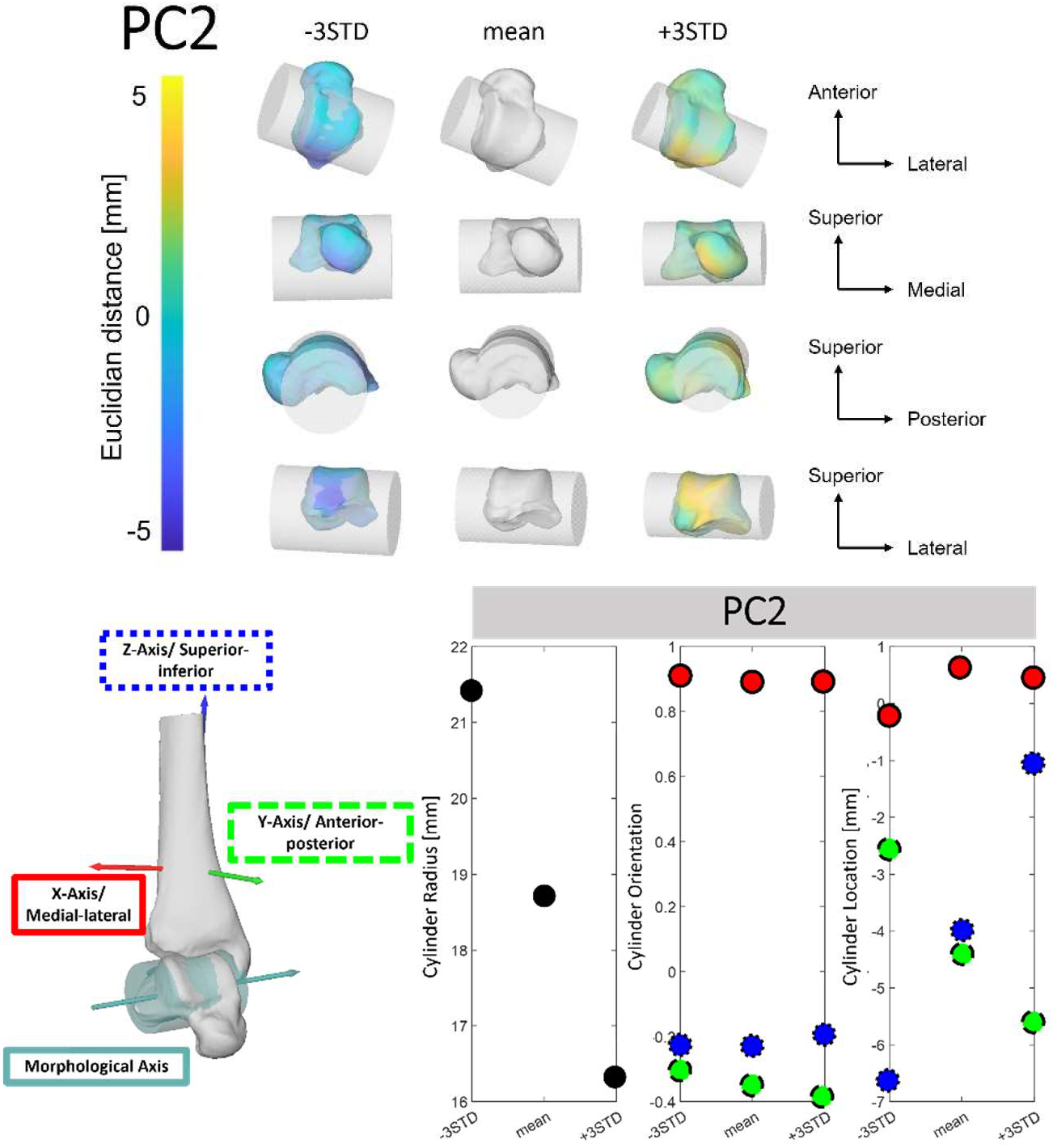
Second PC of talus visualized with cylinder fit through talar dome ± 3 STD from the mean shape. Shape changes are color-coded according to the Euclidean distance in mm to the mean shape. Cylinder metrics for PC2 depicted across ± 3 STD for changes in cylinder radius (black), orientation and location with the three directions: medial-lateral (red and solid outline), anterior-posterior (green and dashed outline) and superior-inferior (blue and dotted outline). n_Participant_ = 36

**Figure 5:**
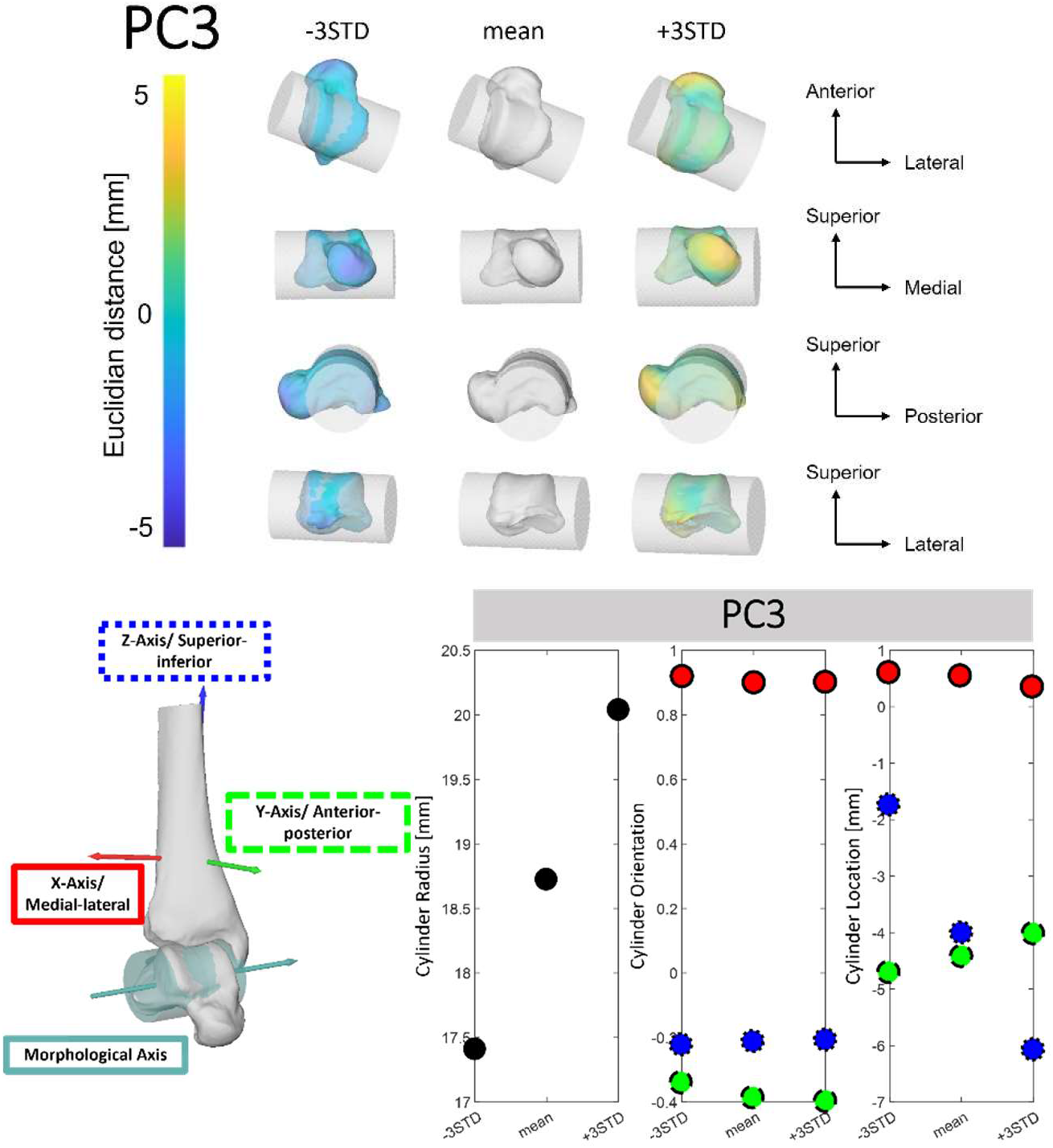
Third PC of talus visualized with cylinder fit through talar dome ± 3 STD from the mean shape. Shape changes are color-coded according to the Euclidean distance in mm to the mean shape. Cylinder metrics for PC3 depicted across ± 3 STD for changes in cylinder radius (black), orientation and location with the three directions: medial-lateral (red and solid outline), anterior-posterior (green and dashed outline) and superior-inferior (blue and dotted outline). n_Participant_ = 36

PC3 explains 11% of the overall variance in the talus. The changes are most prominent in the posterior aspect of the talus and most anterior aspect of the talus head that appear to elongate or shorted the talus. Specially the posterior shape alterations affect the size of the cylinder, hence the talar dome curvature, and the morphological axis location in the superior-inferior direction. For example, the shorter (missing the posterior process) but flatter talar dome (here associated with +3STD changes), the larger the cylinder radius and more inferiorly located the morphological axis. Changes in morphological axis orientations were below 0.1 in all three directions.

Since PC1 of the shape-PCA did not affect the cylinder radius, no regression was performed between PC1 loadings and the cylinder radius. However, regressions for PC2 and PC3 loadings of the shape-PCA and cylinder radius showed significant relationships (Supplement Figure S7). Cylinder radius increased with larger PC2 loadings (p-value = 0.00; r^2^ = 0.47) while the radius decreased with larger PC3 loadings (p-value = 0.0001; r^2^ = 0.09).

## 4. Discussion

This study aimed to quantify the agreement of the morphological (based on a cylinder fit) and rotational axes of the ankle joint, and to determine the morphological features that influence these axes. Overall, we found that the rotation axis was tightly coupled to the morphological axis; however, the morphological axis consistently overestimated the anterior-posterior orientation of the rotation axis. From the shape atlas of the talus, we found that both size and morphological features have independent influences on the curvature of the talar dome and therefore, the location of the morphological axis, as determined by the cylinder fit. The most important morphological features in the shape model showed large variation in the prominence of the posterior section of the talus and the anterior-posterior length of the talar dome next to twisting of the talar head.

Fitting a cylinder to the talar dome served two purposes; first, it allowed us to test the assumption that the talocrural joint acts as a simple hinge (not necessarily only in sagittal plane orientation). Second, it helped simplify the description of the complex shape features captured in our statistical shape model. While the rotation and morphological axes agree well in terms of the axis location, the agreement in orientation was poorer. The morphological axis consistently predicted more talocrural inversion-eversion than was present. This finding indicates that there are aspects of the talus geometry that dictate talus motion which cannot be captured by a simple cylinder. These findings also confirm previous observations [30,31] that the talocrural joint does not behave as a simple hinge, despite the majority of motion occurring in the sagittal plane. While the out-of-sagittal plane motions were small, they contributed to the deviations in the alignment of the morphological and rotation axes. Taking non-sagittal talar dome morphology into account and capturing a greater range of movement, such as cutting maneuvers, might provide a more comprehensive assessment of the non-sagittal relationships between morphology and kinematics.

Despite the small anterior-posterior offset in orientation, the agreement between the axes in terms of location and orientation was sufficient to meet our second goal of using the cylinder fit to help describe how variation in size and shape of the talus alters the morphological axes. We were surprised by how well the cylinder captured function of the talocrural joint, considering that other papers deemed a cylindric shape too simple for the talocrural joint [30,31]. The differences in interpretation might be because these studies were not comparing their morphological axis to the rotation axes. Instead, they assessed how well their geometrical shapes fit the raw bone shape. We suggest that validating more complex shapes used in these studies [30,31] with *in vivo* kinematics may be a suitable next step to further understand the shape-function relationships within the ankle joint.

Talar dome curvature is an important feature for implant design because it may alter the location of the morphological and therefore rotation axis, affecting muscle moment arms, contact pressure, and muscle-tendon and ligament operating ranges. In our herein introduced shape model, we captured talar dome curvature by the radius of the cylinder fit, as well as the vertical location of the morphological axis. Since the morphological and rotation axes’ locations agree well, we assume that changes in the talar dome curvature via size or shape changes are accompanied by alterations in superior-inferior rotation axes location of the talocrural joint. Conceptually, a less curved talar dome corresponds to a larger cylinder, hence lower vertical location of the morphological and rotation axis, resulting in larger moment arms for tendons and muscles attaching to the talus. This would lead to a larger mechanical advantage but also larger tendon excursions for the same angular displacement. Larger tendon excursions result in higher muscle velocities, which lead to reduced forces, according to the force-length relationship [17,18]. Additionally, ligaments and tendon operating ranges may increase, causing the tissues to excessively strain. In muscle mechanics research, this effect is typically referred to as gearing [18]. The opposite behaviour could occur when the talar dome curvature of the implant is too small for the individual (more curved talar dome). We found that overall talus size is strongly linked to talar dome curvature, indicating that larger bones typically have larger talar domes. This makes sense; however, we also found that a substantial amount of the variation in the cylinders’ vertical axis location was explained by shape features that were independent of size. This is particularly interesting for implant design, as the bone might not scale isometrically across all features. While more research is required to confirm the consequences of mismatching talar dome curvature, these findings lend support to the emerging practices of developing person-specific implants.

When interpreting our findings, it is important to consider several limitations. First, we computed our rotation axes over a large range of motion. While this improves the robustness of the helical axis metric, it neglects the nuanced motion that occurs within the range of motion. Since our comparison was to a morphological axis that does not change over the gait cycle, the time series motions would not affect our conclusions, as our rotation axis captures the gross motion of the talocrural joint. Second, BVR data is difficult to track, and we were limited to a small sample size. Rather than directly comparing shape to our rotation axes we used the cylinder axis as a proxy. Future studies with a larger sample size may be able to glean new insights by directly comparing kinematics to specific shape features. Third, since we wanted to explore how talar dome morphology influenced talocrural kinematics, we did not extend our analysis to include the talus facet on the tibia. We made this simplification based on the assumption that the tibia shape would closely mirror the talar dome shape. New insights may also be gained by accounting for the tibia surface and the subsequent joint congruence [1]. Finally, we examined hopping as a model of locomotion (instead of walking or running) to ensure a large range of motion to compute the rotation axis and greater data yield per-participant (more frames of data collected). While we have previously shown the kinematic behaviour of the ankle joint to be similar in running and hopping [9], it will be important to extend future research to running and walking. That said, the talocrural joint is highly congruent and the hopping motion captures its full range of primary motion. Despite these limitations, our approach allowed us to determine how well the morphological axis was able to capture the rotation axis and determine which shape features of the talar dome alter the axes.

## 5. Future Applications

Understanding the relationship between the talocrural joint’s size, shape, and motion is an important progression to advance development and personalisation of effective ankle joint implants for patients requiring ankle joint arthroplasty. Most common talus prostheses differ in overall size, but do not account for individual differences in bone shape, particularly variations in talar dome shape [31]. This might affect post-surgical ankle joint mechanics, prosthesis lifespan and overall quality of life for patients who undergo these procedures [32–34]. If an implant aims to restore “natural” talocrural kinematics, personalising both size and shape features might be an important next step in this process.

## 6. Conclusion

Here we have shown that a cylinder fit to the talar dome provides a reasonable estimate of talar dome curvature. Combining this geometric model with high-resolution BVR data, we have revealed that the axis projecting through the centre of the cylinder (morphological axis) aligns closely with the location, and to a lesser extent, the orientation of the rotation axes of the ankle joint. Despite only small offsets in the orientation of the morphological and rotation axes, we report systematic offsets in ankle joint inversion and eversion motion. Thus, small changes in talus morphology and the associated influence on axis orientation, produce meaningful kinematic alterations that might greatly affect joint contact forces [35]. Future research should investigate how more complex geometric fits perform to replicate talocrural kinematics.

## Notes

Competing Interest AVB, LW, MJR, and LK declare that they have no conflict of interest. AVB was supported by the Matching Dissertation Grant by the International Society of Biomechanics. LK and MJR were supported by the Australian Research Council Discovery Grant (DP160101117). LAK also received funding by the Australian Research Council DECRA (DE200100585). MJR was further supported by the Natural Sciences and Engineering Research Council of Canada Discover Grant (NSERC RGPIN 04880-2022). LW was funded by the Natural Sciences and Engineering Research Council Postdoctoral Fellowship (NSERC PDF 558140-2021).

### Competing Interest Statement

The authors have declared no competing interest.

