## Supplemental Figures for "The influence of talus size and shape on *in vivo* talocrural hopping kinematics"

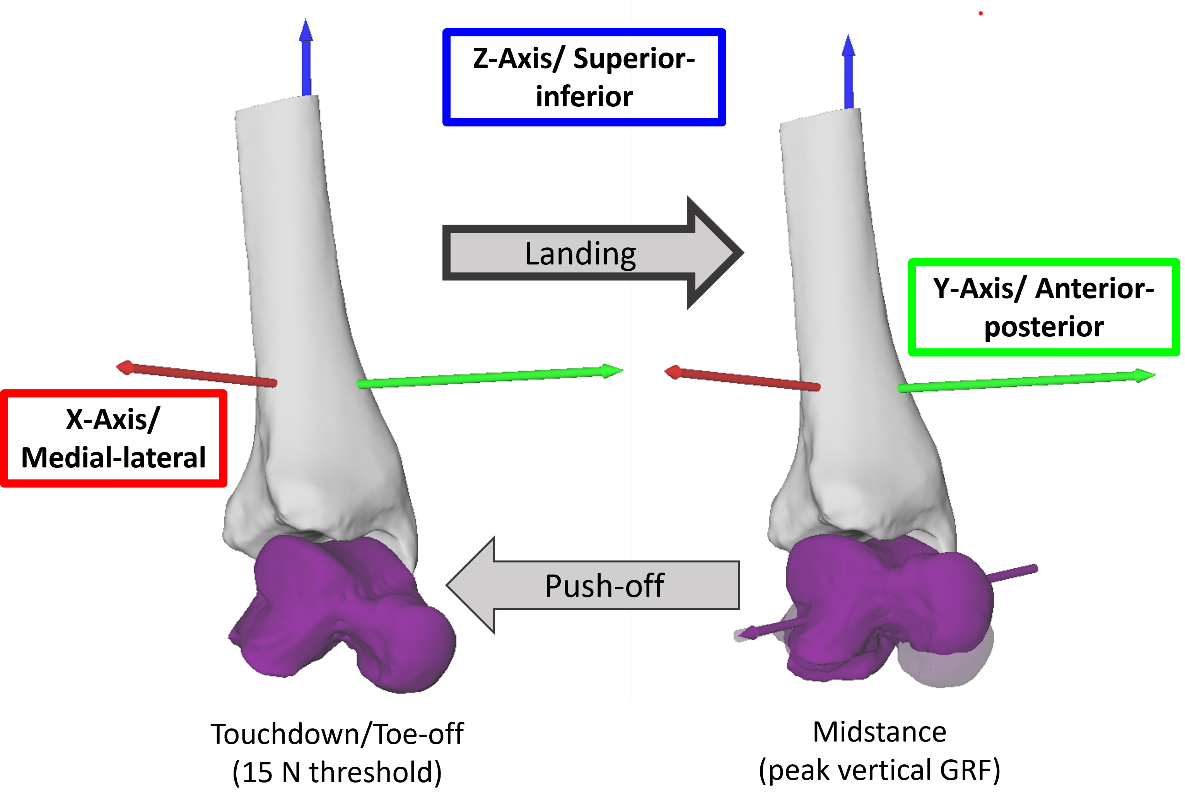

Figure S1: Example of a helical/rotation axis (purple arrow) for landing (n_Participant_ = 1; n_Trials_hopping_ = 3). Talus position at touchdown on the left and during change of position from touchdown (transparent) to the transition point (solid) on the right. The cone about the rotation axis indicates the standard deviation across all three hops. The tibia coordinate system is indicated with red, green, and blue arrows.

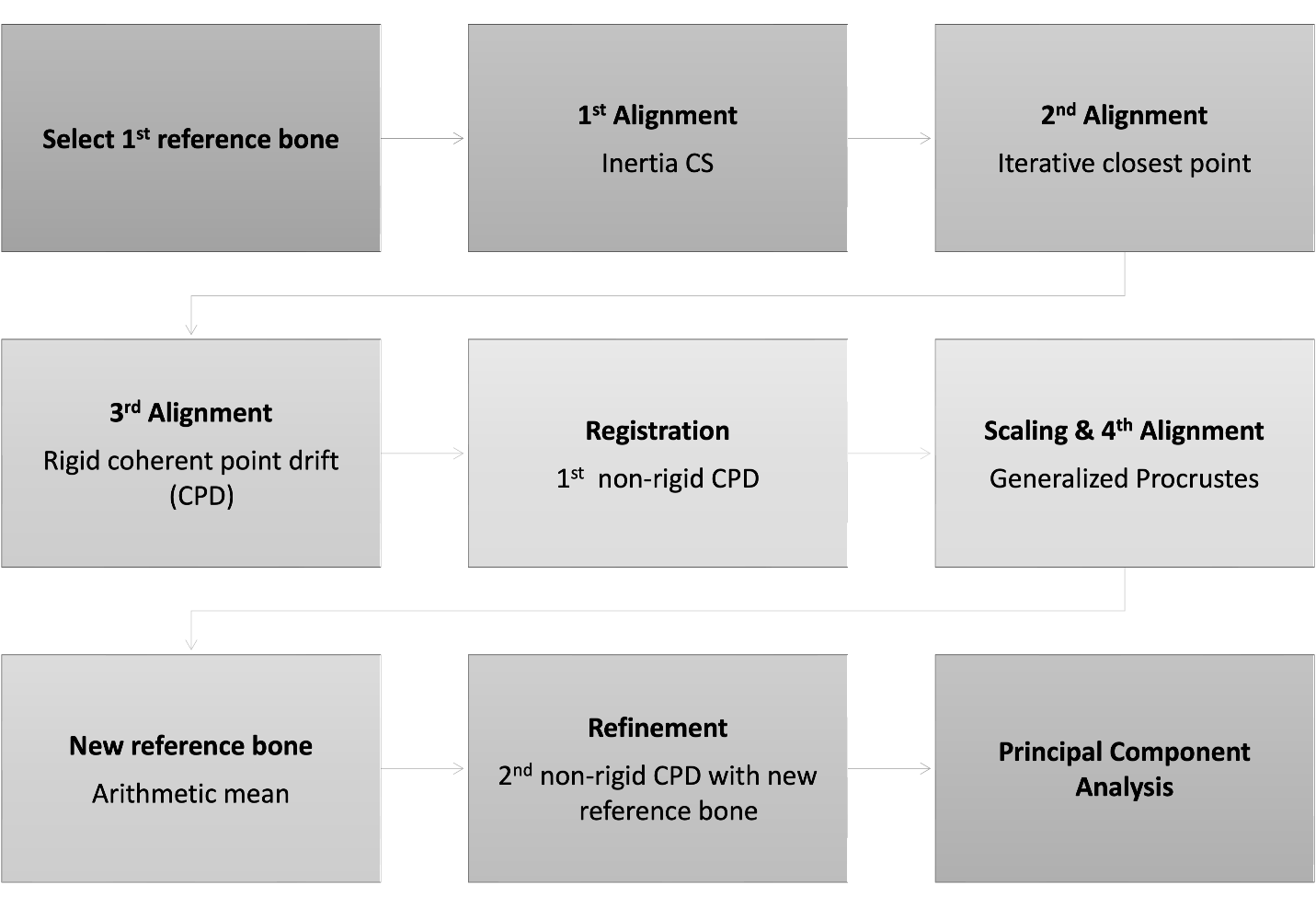

Figure S2: Chronological steps to build shape atlas of 36 tali bones based on the segmented computed tomography scans.

Table S1: Spatial 3D angle between the morphological and rotation axis for each participant and the overall average and standard deviation (STD) for the landing and push-off phase., respectively.

|  | **Landing** | **Push-off** |
| --- | --- | --- |
| **Participant 1** | 11.3 | 10.1 |
| **Participant 2** | 8.7 | 12.1 |
| **Participant 3** | 6.4 | 6.4 |
| **Participant 4** | 10.9 | 13.8 |
| **Participant 5** | 14.0 | 12.0 |
| **Participant 6** | 5.6 | 3.6 |
| **Participant 7** | 2.5 | 6.1 |
| **Participant 8** | 10.8 | 11.2 |
| **Participant 9** | 10.5 | 11.3 |
| **Mean ± STD** | 9.0 ± 3.5 | 9.6 ± 3.4 |

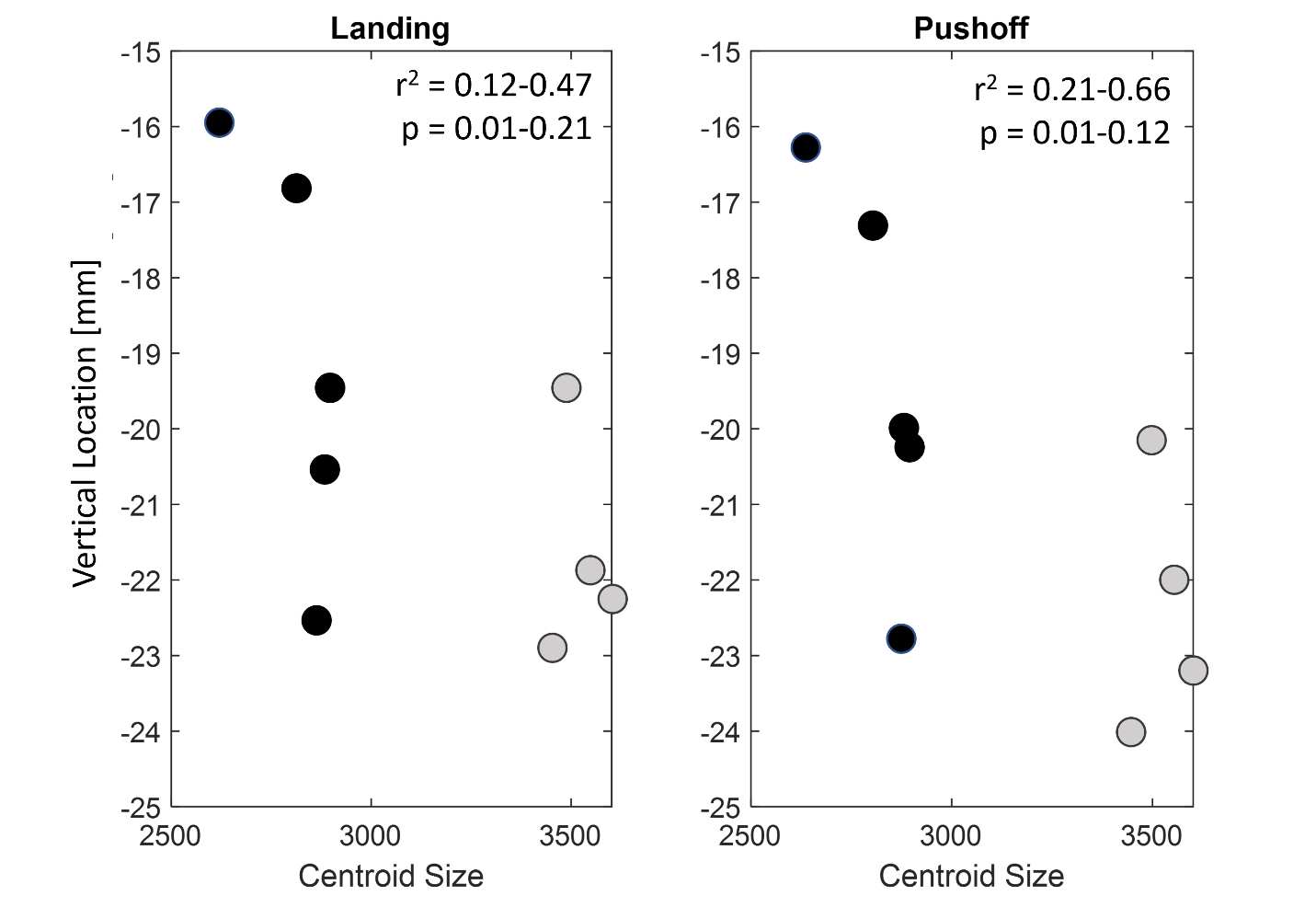

Figure S3: Linear regression of vertical location of the rotation axis and centroid size (n_Participant_ = 9) during single leg hopping. Females are indicated as gray circles while males are black. P-values and coefficients of determination were calculated with the leave-one-out method and provided as a range from minimum to maximum.

Figure S4: Orientation and location of rotation (average across all trials; purple color) and morphological (teal color) axes for each participant during push-off.

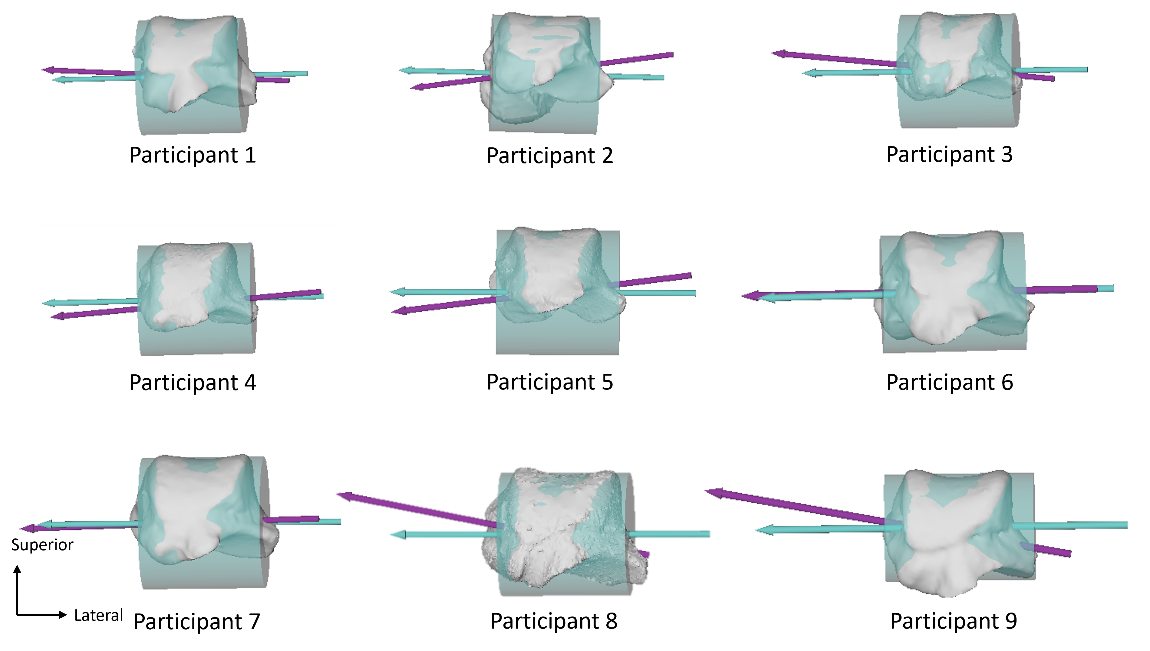

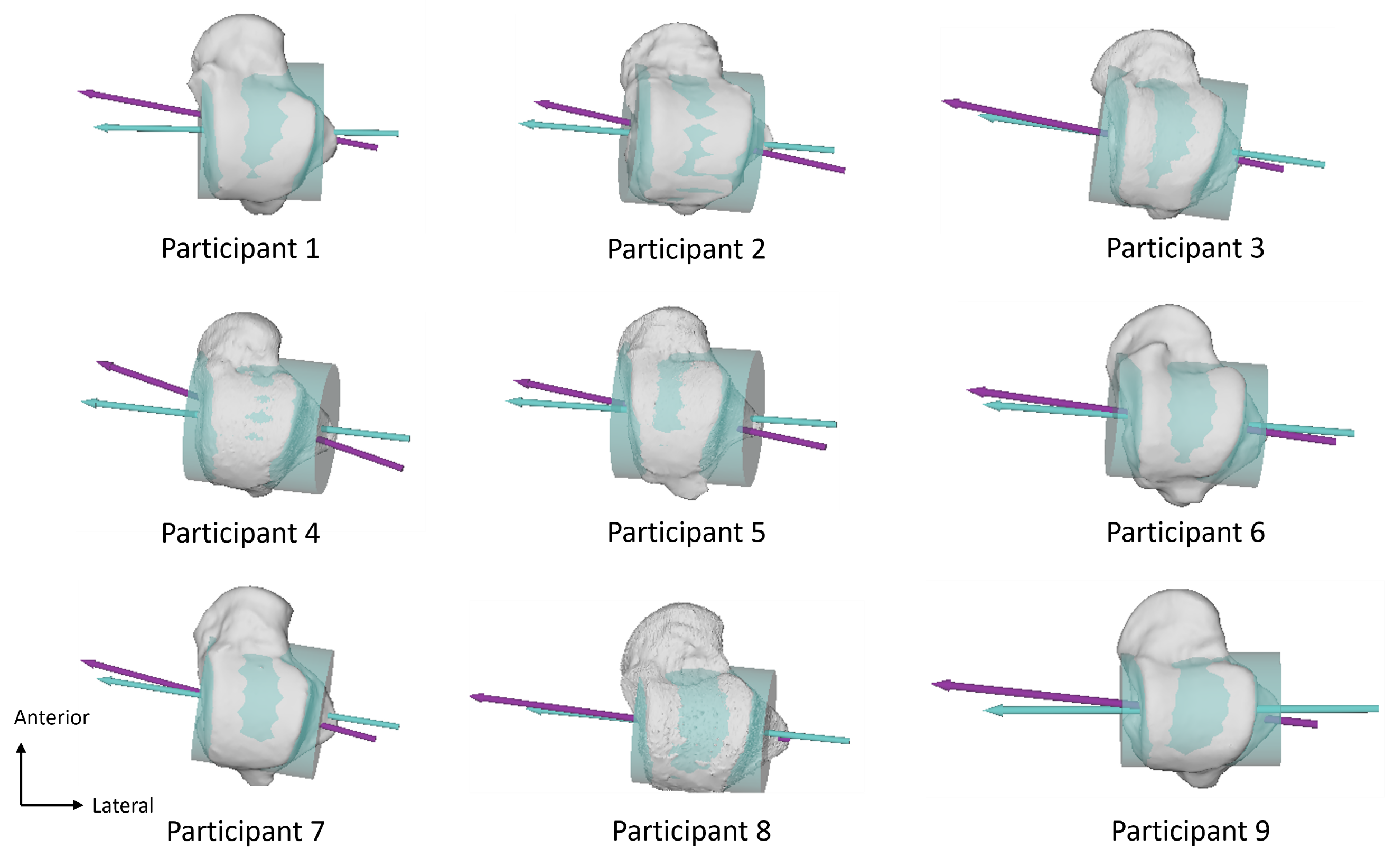

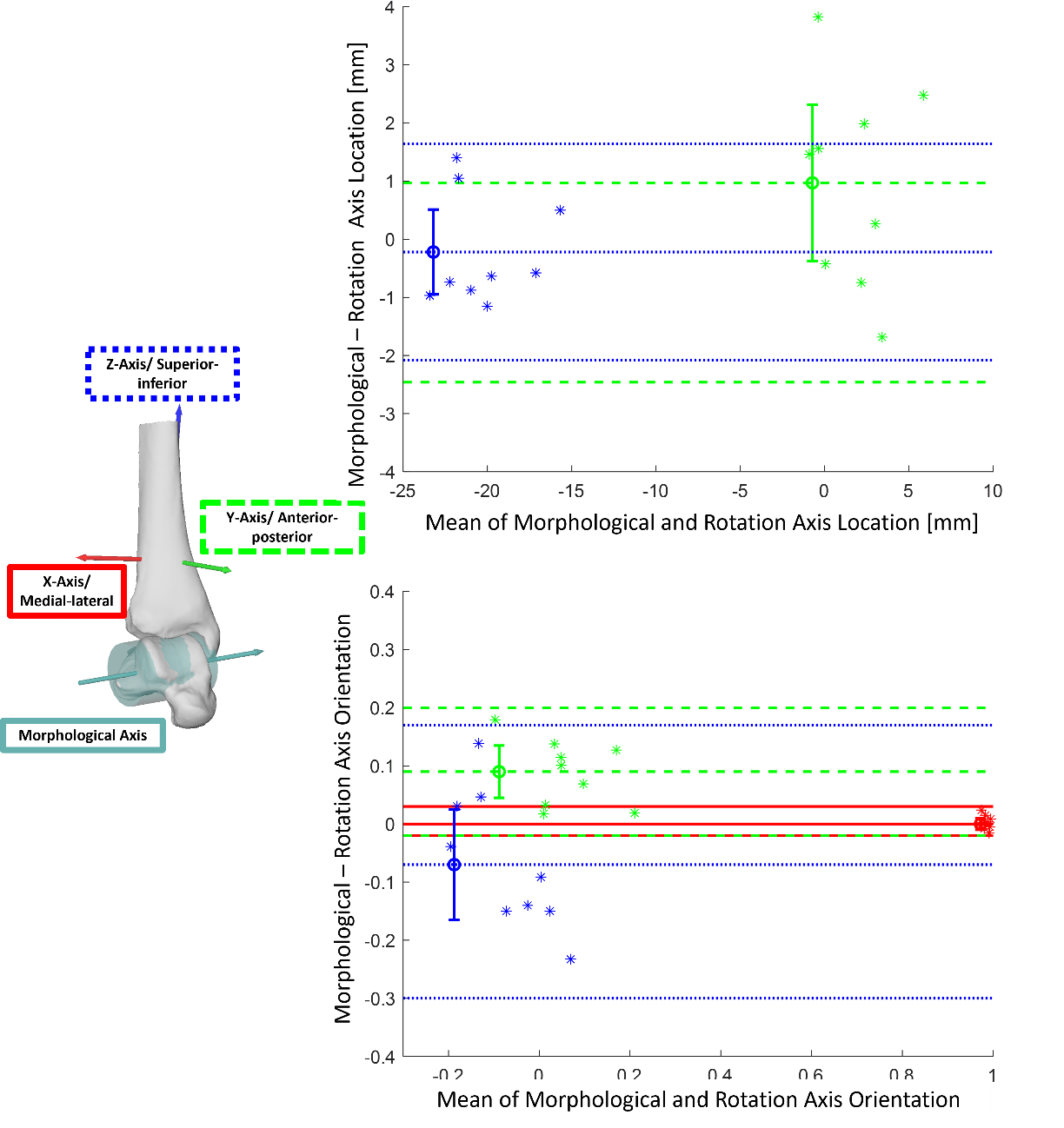

Figure S5: Bland-Altman plots indicating level of agreement between the morphological and rotation axis location (top) and orientation (bottom) during landing. Directions are color-coded according to the legend on the side. The line with the error bars indicates the overall mean ± 95% confidence interval and the other two lines indicate the limits of agreement. Note, there is not medial-lateral values for the location as the helical axis location was evaluated based on where they penetrate the sagittal plane of the tibia coordinate system.

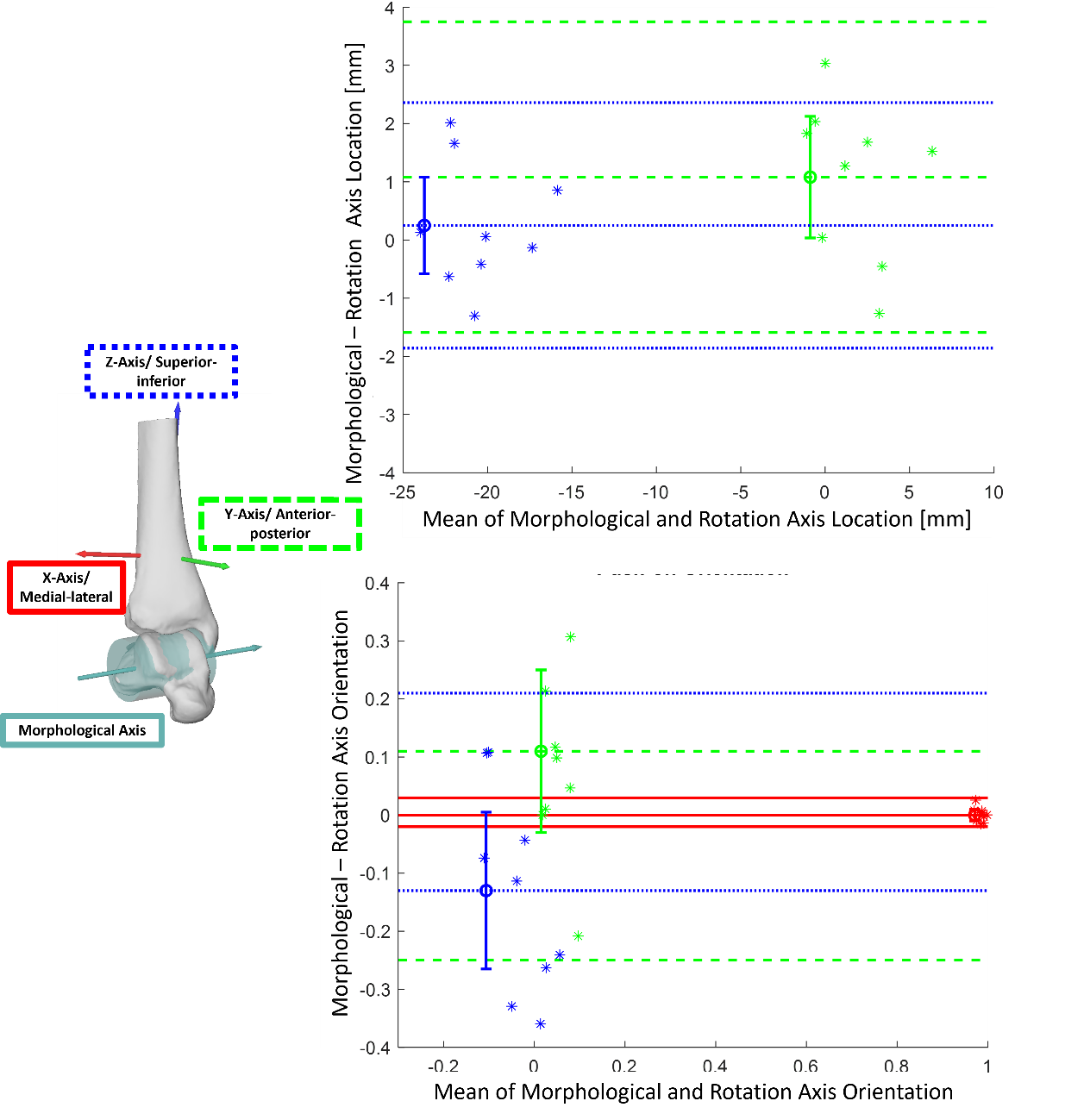

Figure S6: Bland-Altman plots indicating level of agreement between the morphological and kinematic axis location (top) and orientation (bottom) during push-off. Directions are color-coded according to the legend on the side. The line with the error bars indicates the overall mean ± 95% confidence interval and the other two lines indicate the limits of agreement. Note, there is not medial-lateral values for the location as the helical axis location was evaluated based on where they penetrate the sagittal plane of the tibia coordinate system.

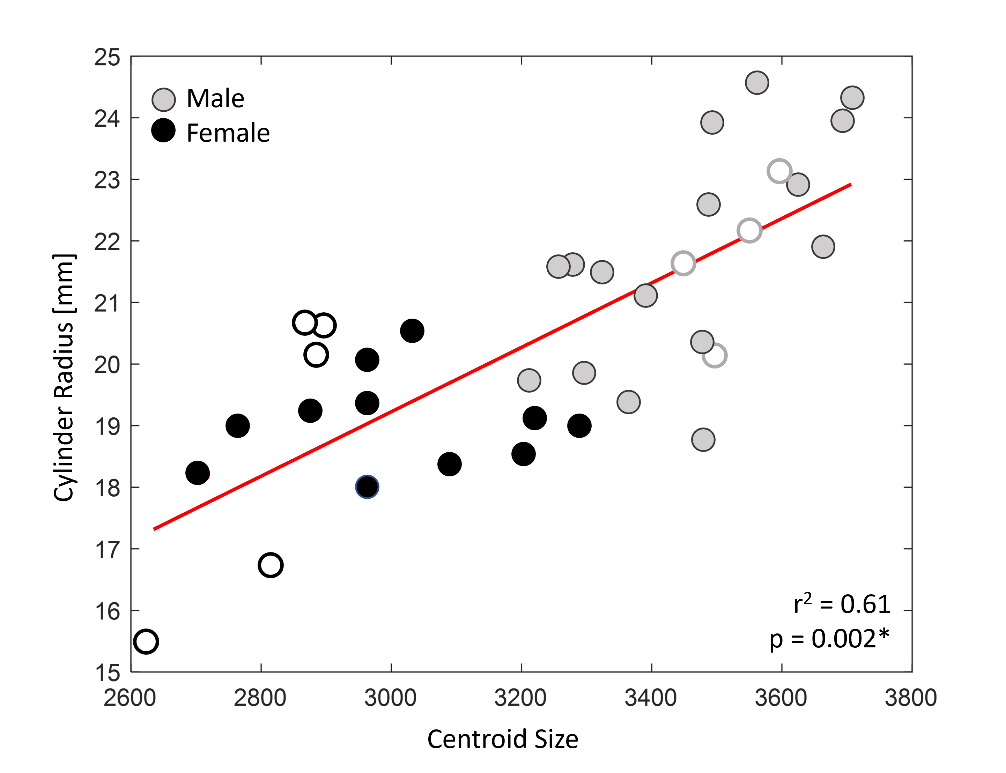

Figure S7: Linear regression of cylinder radius on centroid size (n_Participant_ = 36). Grey circles indicate males while black circles indicate females. Empty circles indicate participants with hopping data.

9

Figure S8: Linear regression (red line) of the cylinder radius on shape loadings of PC2 (top) and PC3 (bottom). Grey circles indicate males, black circles females. n_Participant_ = 36.

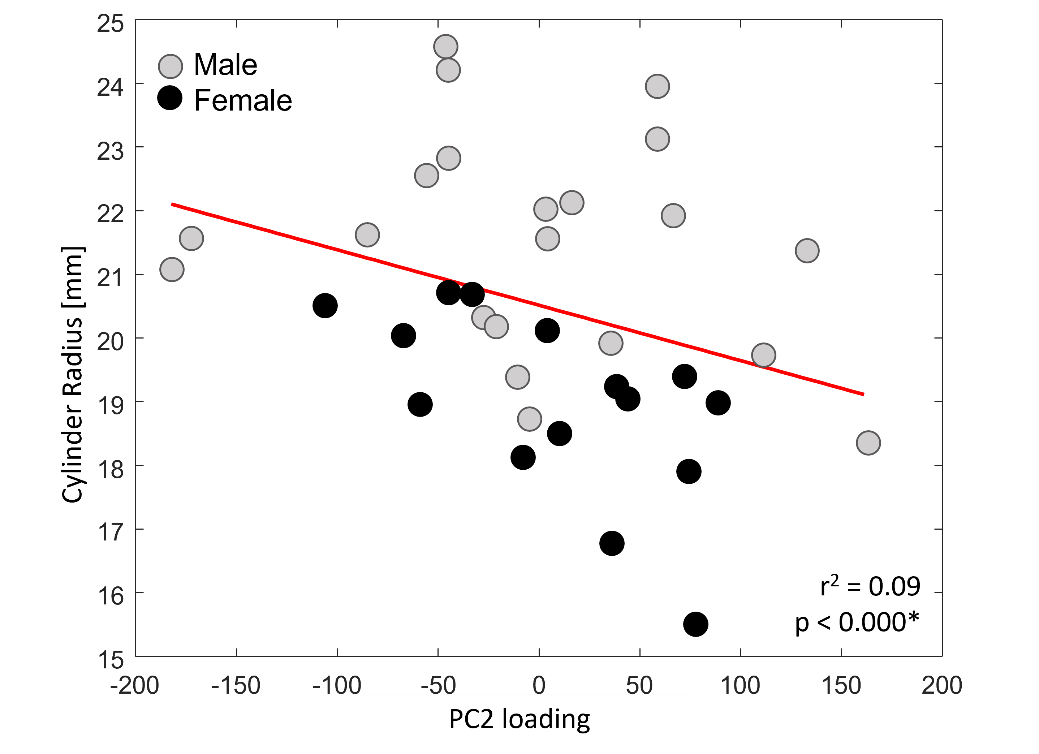

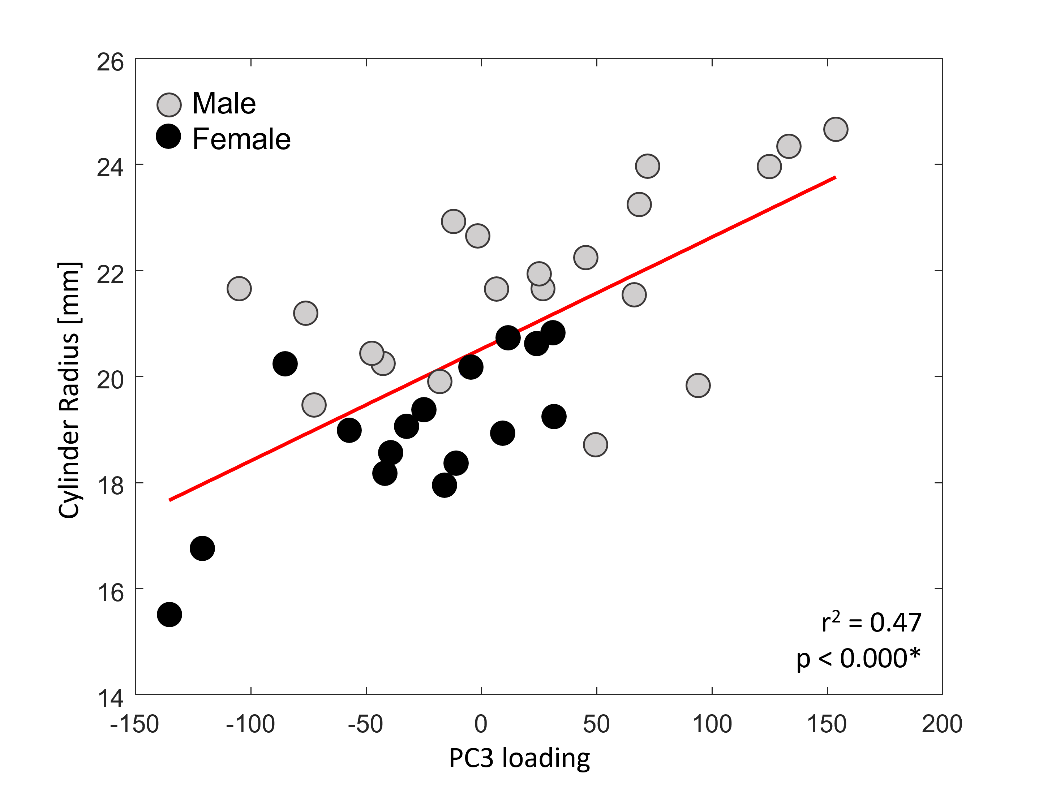

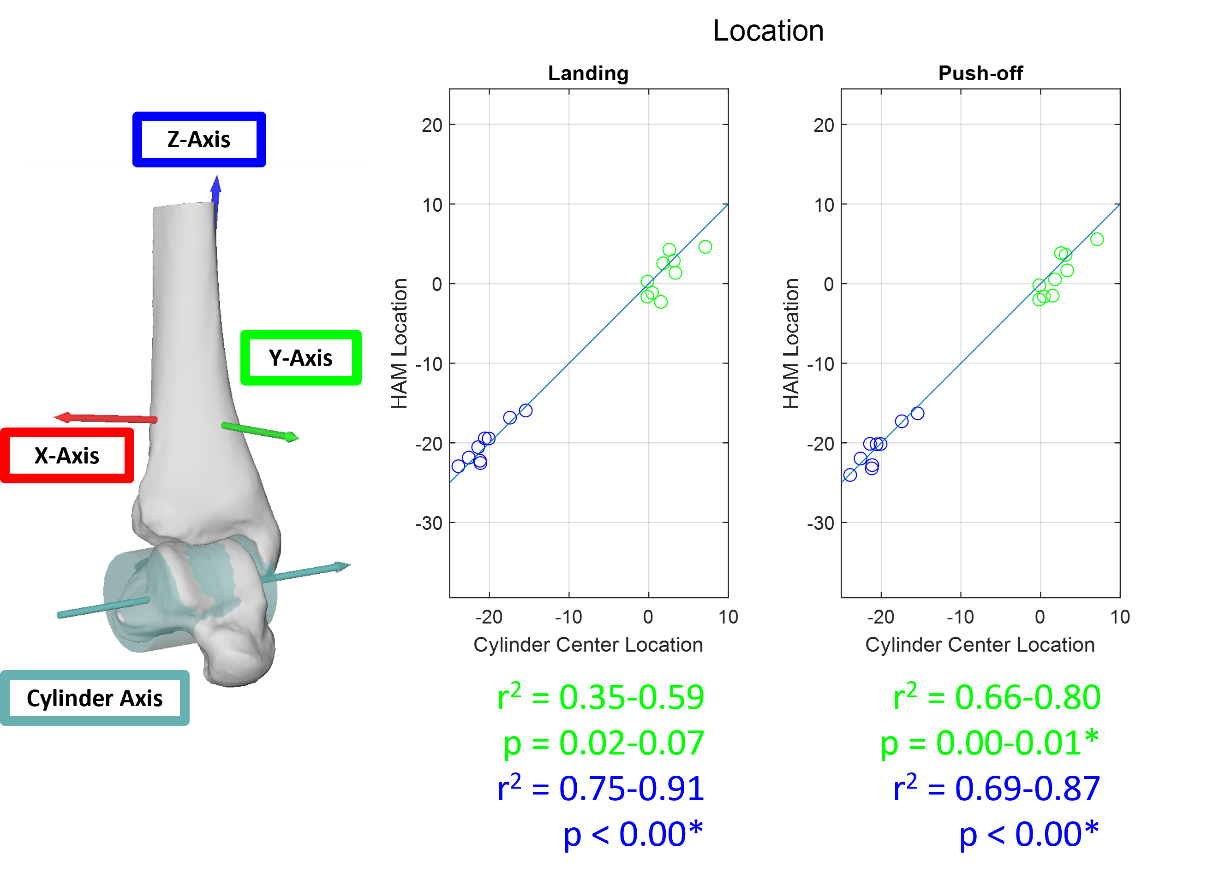

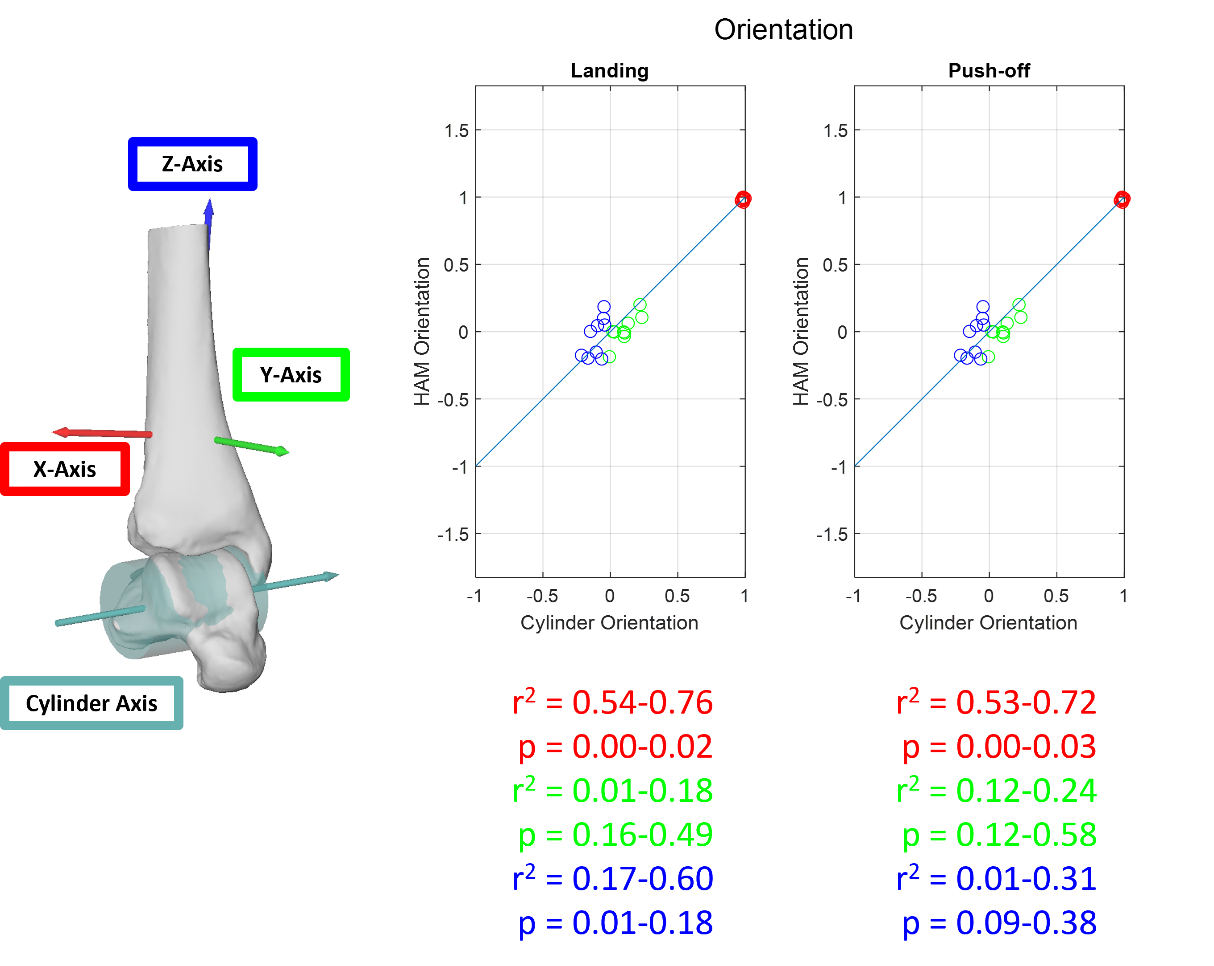
